# Eye tracking insights into movement preparation and execution under nonstandard visual movement feedback

**DOI:** 10.64898/2026.02.04.703867

**Authors:** Felix Quirmbach, Jens R. Helmert, Sebastian Pannasch, Annika Dix, Jakub Limanowski

## Abstract

For eye-hand coordination, predictions of sensory movement consequences may already be issued, and adjusted, during action preparation. In this pre-registered study, we combined a delayed-movement paradigm with a virtual reality-based hand-eye tracking task to investigate the oculomotor correlates of planning and executing coordinated hand-eye movements under standard vs nonstandard visual hand movement feedback. We measured pupil dilation and gaze-hand tracking during action preparation and subsequent task execution, where visual movement feedback violated or matched cued expectations: Participants prepared and, after a delay period, executed hand movements. Their movements were reflected by congruent or incongruent (inverted) movements of a glove-controlled virtual hand model, which they had to follow with their gaze. In the preceding delay period, visual cues could specify the to-be-executed movement (or leave it unspecified), and the visuomotor mapping (congruent or incongruent, 75% cue validity). We found that during the delay, pupil diameter increased more strongly when the movement was pre-cued (compared to left unspecified), and when nonstandard compared to standard visual movement feedback was expected. During execution, gaze-hand tracking performance decreased under nonstandard mappings, but significantly less so when the to-be-executed movement was pre-cued. Expectation violation trials produced a strong pupil dilation, particularly when congruent (standard) visuomotor expectations were violated, but also when incongruent mappings were cued but congruent ones observed. Furthermore, expectation violation impaired tracking performance; again, stronger for pre-cued movements with standard mapping. Our results indicate that oculomotor responses during delay encode processes related to motor planning and flexible forward prediction of sensory action consequences ahead of execution, i.e. increased mental effort and expectations of sensory conflict. Moreover, the results demonstrate that the strength of these (updated) predictions affects eye-hand coordination and pupillary responses during subsequent execution of the planned action.

## Introduction

Performing everyday manual actions like grasping an object or catching a ball requires the coordinated control of hand and eye movements. This often involves tracking the hand’s movements with one’s gaze. Thus, besides coordinating the motor commands issued to the different effectors, it requires an accurate prediction of where the hand will be seen and what the hand movement will look like. In other words, it relies on an adequate prediction of the sensory data generated by the (hand) movement. The prediction of sensory action consequences is thought to be enabled by internal forward models in the brain (Miall & Wolpert, 1996; Shadmehr & Krakauer, 2008; Wolpert & Kawato, 1998). These forward models are based on life-long learning, yet they are also surprisingly flexible. For instance, participants can quickly adapt to novel sensorimotor mappings, such as when learning to move under nonstandard (e.g., rotated or delayed) visual movement feedback (e.g., Bock, 2013; Cunningham, 1989; Limanowski et al., 2017). This learning process happens through updating forward models by sensory prediction errors; i.e., unpredicted sensory action consequences (Grafton et al., 2008; Shadmehr et al., 2010; Synofzik et al., 2006; Tseng et al., 2007). Notably, the concept of forward sensory predictions extends to action preparation; that is, generating sensory predictions during motor planning—even in the absence of actual movement and related sensory feedback (Kilteni et al., 2018; Kuang et al., 2015; Sawtell, 2017; van Kemenade et al., 2016). Indeed, recent brain imaging and electrophysiological work using delayed-movement paradigms has shown that the preparation of (delayed) actions under nonstandard visual movement feedback is associated with increased activity in brain regions thought to implement forward models; i.e., the cerebellum and posterior parietal cortex (Kuang et al., 2015; Pilacinski et al., 2018; Quirmbach & Limanowski, 2024). This implies that the brain’s forward models can anticipate novel (“unpredicted”) sensory action consequences, and adjust their respective sensory predictions ahead of execution.

Besides central nervous activity, oculomotor behavior can reveal much about the mental processes associated with sensorimotor integration and the related predictive processes for eye-hand coordination. Particularly promising is pupil dilation, as a distinct readout of locus coeruleus-noradrenergic responses related to executive control (Alnæs et al., 2014; Grujic et al., 2024). Thus, pupil dilation is considered a reliable marker of the (task-relevant) uncertainty of internal states (Becker et al., 2024; Fan et al., 2023; Preuschoff, 2011; Satterthwaite, 2007), and of mental effort (Aston-Jones & Cohen, 2005; Joshi & Gold, 2020; Shenhav et al., 2017), also in sensorimotor tasks (Hosseini, 2017; cf. Zénon, 2014). Pupil dilation increases when responses are pre-cued (Adam et al., 2014; Moresi et al., 2008), and pupil diameter can reflect the anticipated difficulty or the expected reward of a task (Dix & Li, 2020; Irons et al., 2017; Spliethoff et al., 2022). Furthermore, expectation violation reliably evokes pupil dilation (Alamia et al., 2019; Bianco, 2020; Braem et al., 2015; Kloosterman et al., 2015; Reisenzein et al., 2019; Yokoi & Weiler, 2022a), and error-based motor learning is reflected by changes in trial-by-trial pupil dilation (Yokoi & Weiler, 2022a). This suggests a possible link of pupillary responses to the updating of sensorimotor associations and the sensory predictions of internal (forward) models. Specifically, pupil size could indicate the mental effort and the (un)certainty associated with generating (nonstandard) sensory forward predictions.

Pupil size has been measured to assess anticipatory mental effort during saccade preparation in a delayed movement task, but without concurrent hand movements (Hutchison et al., 2020; Jainta, 2011; Unsworth et al., 2023; Wang, 2016; Wang et al., 2015). Several eye tracking studies have used anti-saccade designs to decouple hand and eye movement goals; i.e., requiring the execution of eye and hand movements to different (vs the same) targets or directions. These studies have yielded important insights into the on-line coordination of hand and eye movements under various conditions, including saccadic vs smooth pursuit, or manipulated visual movement feedback (e.g., (Chen et al., 2016b; Gorbet & Sergio, 2009; Vercher et al., 1995; M. Yeomans et al., 2021). However, these studies focused on on-line control; i.e., execution without prior planning or preparation. Thus, to our knowledge, no one has yet investigated oculomotor behavior during the preparation (and subsequent execution) of manual actions with nonstandard visual hand movement feedback.

Therefore, in this pre-registered study (https://doi.org/10.17605/OSF.IO/6KM9Y), we built upon the above works by investigating the oculomotor correlates of preparing and executing manual movements under nonstandard visual hand movement feedback, under varying certainty of the underlying movement plans. For this, we employed a novel hand-eye tracking task (Fig. 1): Participants controlled a photorealistic virtual hand via a data glove worn on their unseen hand, while tracking the motion of the virtual hand with their gaze. Crucially, we embedded this task in a delayed-movement paradigm; i.e., participants prepared the to-be-executed hand movements and executed them after a delay. During the delay (preparation period), visual cues could specify the to-be-executed movement and the expected visuomotor mapping. Specifically, the cue shape could indicate the preparation of an open or a close hand movement (or leave it unspecified), while the cue color indicated whether, in the upcoming execution period, the virtual hand would display the participant’s actual executed hand movement (congruent mapping) or an inverted movement (incongruent mapping). Importantly, the virtual hand movements were always controlled by the participant; i.e., in the incongruent trials, the virtual hand received the same glove sensor values, only inverted. Thus, for efficient gaze-hand tracking, participants needed to generate the appropriate visuomotor predictions; i.e., movement plans and predictions of visual hand movement feedback. In the incongruent conditions, the predictions of how the executed hand movement will look like differed from the standard (life-long learned) association and, consequently, required updating for subsequent gaze-hand tracking. Finally, we added “expectation violation” trials, in which the virtual hand moved contrary to the cued visuomotor mapping, and thus, the visual movement feedback violated the participants’ expectations. During the task, we recorded hand and gaze movements, as well as pupil diameter. We hypothesized that these measures would indicate the mental effort involved in anticipating conflicting (nonstandard) visual movement feedback and updating visuomotor expectations during planning; and how these factors influenced gaze-hand coordination during subsequent task execution. Specifically, we had six key hypotheses:

**Figure 1:**
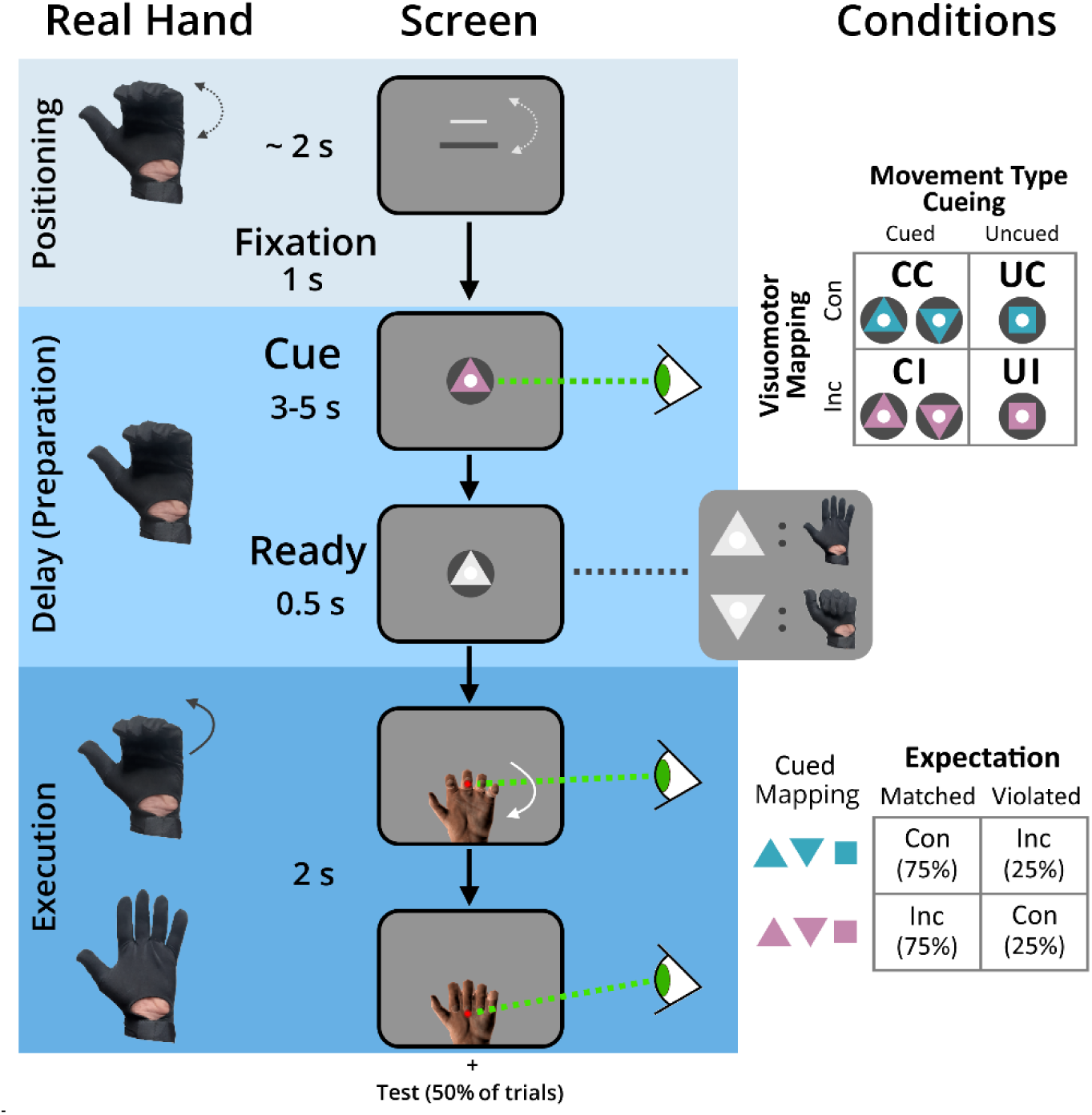
Task Design. Schematic of an exemplary trial. The participants’ unseen real hand movements were measured via data glove (left) and fed to a virtual hand model presented on screen (middle); with visual cues (right) predicting different aspects of the upcoming, to-be-executed movement. To begin a trial, participants moved their hand to a neutral, half-opened starting position, by aligning a moving bar with a target (see Methods). After a 1 s fixation-only period, a visual cue was then presented for 3-5 s. The cue color (cyan or pink) indicated the visuomotor mapping during subsequent movement execution: In 50% of trials, the virtual hand movement would reflect the movements of the real hand (congruent condition, here: cyan), in the other 50% its movement was inverted (incongruent condition, here: pink). The cue shape could predict the to-be-executed hand movement: either opening (upward pointing triangle, 25%) or closing it (downward triangle, 25%). In the remaining 50% of trials, a square cue shape left the to-be-executed movement unspecified. After the delay period, a brief (0.5 s) ‘Ready’ signal (white triangle) appeared, indicating that movement execution would follow immediately and instructed subjects to either open (triangle pointing up) or close (triangle pointing down) their real hand—in cued trials, this would confirm the previous movement type, while signaling it for the first time in uncued trials. Subjects then had to execute the instructed hand movement during a 2 s time period following the appearance of the virtual hand. During hand movement execution, participants had to follow a red dot at the middle finger tip of the virtual hand with their gaze. As cued during delay, the virtual hand moved either congruently or incongruently (i.e., inverted) to the real hand. Importantly, the mapping cue was only valid in 75% of trials. In the remaining 25% of trials, the virtual hand moved opposite to what was cued (e.g., congruently, although an incongruent mapping had been cued), to violate cued visuomotor expectations. Overall, this resulted in a full factorial, within-subject design with factors *Movement type cueing* (Cued, Uncued) and *Visuomotor mapping* (Congruent, Incongruent), and additional nested factors specifying *Movement type* (Open, Close) and *Expectation violation* (Matched, Violated). Stimuli and virtual display not to scale.

During the delay phase (movement preparation), we expected greater pupil dilation to be evoked by cueing incongruent>congruent visual movement feedback (main effect, *Hypothesis H1*), reflecting the anticipation of visual hand movement feedback conflicting with life-long visuomotor expectations, the potential updating of forward sensory predictions, and the mental effort associated with correspondingly increased cognitive control (cf. Shenhav et al., 2017; Yon, 2018). We also expected increased pupil dilation if the upcoming movement was cued>uncued (main effect, *Hypothesis H2*). This prediction followed from our previous study using a similar task design (Quirmbach & Limanowski, 2024; cf. Rosenbaum, 1980), where the movement data clearly suggested that participants prepared a specific movement if cued correspondingly, but abstained from (parallel) movement planning in the uncued condition. Furthermore, we expected an interaction effect; i.e., the planning-related (cued>uncued) pupillary dilation should be more pronounced in trials with incongruent compared to congruent visual feedback (*Hypothesis H3*).

During execution, we expected that a successful generation (and, for cued incongruent mappings, an updating) of visuomotor expectations during movement preparation would be associated with a “surprise response” to observing a visual movement contrary to that predicted by the cue (*Hypothesis H4*). This response should be reflected by larger pupil dilation and reduced gaze-hand tracking accuracy in expectation violation trials (see above). Furthermore, we expected better task (oculomotor) performance (less gaze-hand tracking error and faster initiation of hand and eye movements) for movements under congruent, compared to incongruent visuomotor mappings, as the former correspond to lifelong associations between motor commands and visual feedback (*Hypothesis H5*). Previous work has shown that eye tracking of a target controlled by one’s hand movements worsens significantly when the target’s movement is manipulated with respect to the actually executed movement (Chen et al., 2016b; Vercher et al., 1995a; M. Yeomans et al., 2021). Thus, despite potential anticipatory processes and updated expectations, we assumed a residual benefit of moving under standard vs nonstandard visual feedback. Finally, we expected improved task performance when the movement type was cued, compared to being left uncued in the preparation period, because a more specific movement plan could be generated during the respective delay (*Hypothesis H6*).

## Materials and Methods

### Participants

40 healthy volunteers (23 female, mean age = 25.9 years, range 19-36, normal or corrected-to-normal vision) participated in the experiment after signing informed consent. Based on a recent study using a similar task design (Quirmbach & Limanowski, 2024), we determined a minimum sample size of 36 participants to reach 95% power via a-priori power analysis with the software program G*power (Cohen’s *f* = 0.25, alpha error: 0.05, 4 measurements per condition, correlation of 0.5 between measurements). One subject had to be excluded due to technical issues with their eye tracking measurements, leaving 39 participants’ data sets for the group level analysis. As compensation for performing the study, participants received 20 €, and up to 5 € depending on the percentage of catch trial questions answered correctly (1 € if they reached at least a 75% correct answer rate, 3 € for at least 85% correct and 5 € for at least 95% correct, see below). The experiment was approved by the ethics committee of the Technische Universität Dresden and conducted in accordance with this approval.

### Pre-registration

This study was pre-registered (osf.io/6km9y). Preregistration was done after data collection but before data analysis; that is, aside from confirming basic file readability, the content of data files was neither examined nor analyzed in any form prior to registration. Hypotheses, task design and data analyses presented here follow the plan laid out by pre-registration; all deviations from our original plan and exploratory analyses are clearly stated and explained in the article.

### Experimental design and procedure

Participants sat in a constantly lit room on a desk with their heads stabilized via a chin rest, their eyes distanced 93 cm from a screen (BenQ LCD XL2420T), resulting in a maximal viewing angle of 31.9 degrees (horizontal) and 17.7 degrees (vertical). On the screen, all visual stimuli, i.e., instructions, visual cues and the virtual hand model, were displayed with a spatial resolution of 1920 x 1080 pixels, at a refresh rate of 60 Hz. The virtual display was created with the Blender 3D graphics software package (https://www.blender.org/, Version 2.79) and its Python programming interface.

During the experiment, pupil size and gaze position were recorded monocularly (left eye) at 1000 Hz via a desktop-mounted eye tracking system (EyeLink 1000 Plus, SR Research); a 9-point calibration and validation procedure was performed at the start of each block. A data glove (5DT Data Glove 14 Ultra MRI, 60 Hz sampling rate, full-speed USB 1.1 connection) placed on the participants’ left hand measured each finger’s flexion via seven sewn-in optical sensors (two for each finger; one excluded due to technical issues). For each participant, data transformation from the glove was calibrated carefully prior to the actual experiment based on the possible range of flexion to ensure that movement of the real and virtual hand matched. After averaging across sensors to ensure smooth and coherent visual motion (cf. Limanowski & Friston, 2020), its data was fed to a photo-realistic virtual hand model shown on the screen, which participants were therefore able to control. By placing their gloved real hand under the table and restricting their head towards the screen, throughout the experiment, they received visual feedback only from the virtual hand. The experimenter continuously observed both the virtual display and the participant from a distance, ensuring task compliance.

Figure 1 shows an example trial, described in the following: At the beginning of each trial (hand positioning), participants had to bring their real, unseen hand to the neutral starting position (i.e., fingers half-way closed). A small white bar presented on screen, whose vertical position was controlled via data glove by the real hand’s posture, indicated to which degree the hand was opened or closed. To start the trial, this bar had to align with a fixed, central grey bar (see Fig.1); on average, this took 1.98 s (SD = 0.54 s). Once both bars were aligned, they were replaced by a central white fixation dot (0.310 x 0.310 degrees of visual angle). Participants were instructed to keep their gaze on this dot throughout the entirety of the delay period.

After 1 s, a visual cue was presented centrally and remained visible throughout the delay period (3 to 5 s, jittered & pseudo-randomized, evenly distributed across conditions). This cue provided two types of predictive information: In half of all trials, triangle-shaped cues (width: 1.057 / height: 0.870 degrees of visual angle) informed participants in advance which real hand movement they would have to perform, with an upward-pointing triangle (64 trials, 25%) instructing them to open their hand, while a downward-pointing triangle (64 trials, 25%) instructed them to close it. In the other 50% (128 trials), a square-shaped cue (0.682 x 0.682 degrees of visual angle) did not provide any information about the to-be-performed movement. Therefore, participants were able to generate specific predictions about the sensory feedback of the upcoming movement in only half of the trials. Cue shapes were designed such that their on-screen area was identical and were presented at the same position. Simultaneously, cue color indicated if during execution the virtual hand would move congruently to the real hand (i.e. perform the same movement type), or incongruently (i.e., perform the opposite movement type). In the latter case recorded hand movement data was inverted, so that opening the real hand would close the virtual hand, and vice versa. This way, we ensured that even with incongruent visuomotor mappings, visual feedback was still completely predictable from motor commands, and participants would perceive the visual hand movements as caused by their own actions. Cue colors (cyan and pink) were selected to be near-isoluminant; the color coding was balanced across participants.

Following the delay period, a centrally presented white triangle (width: 1.057 / height: 0.870 degrees of visual angle, same as the cue) served as a ‘Ready’ signal for 500 ms, indicating the imminent execution phase. For trials with square-shaped cues—when movement type was left ambiguous—the triangle would also indicate which movement to perform, with an upward-pointing triangle instructing opening of the real hand, and a downward-pointing triangle instructing them to close the hand. To ensure visual feedback of the entire movement, participants were instructed to only start their hand movement as soon as the ‘Ready’ signal and fixation dot disappeared, and the virtual hand model appeared instead. In the following execution period (2 s), participants had to perform the instructed hand movement while following a visual target (red dot) at the tip of the virtual hand’s middle finger with their gaze as closely as possible. At the start of the execution phase, the virtual hand was displayed in a neutral starting position, so that the red dot was positioned exactly in the middle of the screen. The virtual display was furthermore designed to ensure that both movement types were equidistant when the hand movements were executed completely, i.e., the gaze target would cover the same Euclidean distance on the screen. After finishing the hand movement, i.e., completely opening or closing it, the hand was then to be kept in the final position until the end of the trial. After the execution period ended, the virtual hand model disappeared, and a blank screen was presented for 2 s, marking an inter-trial interval.

In the execution period, we included occasional expectation violation trials (25%, i.e. 64 trials, balanced across conditions). In those trials, the virtual hand displayed the opposite visuomotor mapping to that predicted by the cue color during the delay period; i.e., moving congruently when an incongruent mapping was cued, and vice versa. The inclusion of these trials allowed us to examine the oculomotor effects of expectation violation, testing our Hypothesis 4.

Altogether, this study employed a full factorial, within-subjects design with two factors: *Predicted visuomotor mapping* and *movement type cueing*, resulting in four main conditions: CC (Cued movement type, expecting congruent mapping), CI (Cued movement type, expecting incongruent movement), UC (Movement type not cued, expecting congruent mapping) and UI (Movement type not cued, expecting incongruent mapping). For analysis of both pupil size and tracking performance during movement execution, we also included the nested factors *movement type* (Hand opening, closing) and *visuomotor expectation violation* (Matched, Violated). Participants performed the task under all conditions, with equal amounts of trials with hand opening or closing, movement type being cued or uncued, and congruent or incongruent virtual hand behavior. Visuomotor cues were valid in 75% of trials (192 trials) and invalid in 25% (64 trials).

Finally, to ensure that participants were paying attention to the predictive cues (and therefore able to generate visuomotor predictions), we included “catch trials”, in which participants had to indicate if the mapping of the virtual hand matched the cue’s predictions (cf. Yon et al., 2018). After all invalidly cued trials, and a matching number of validly cued ones, a probe question (“Prediction correct?”) was presented centrally, alongside two answer options (“Yes” / “No”) on the lower left / right side of the screen, which could be answered via button press with the right (i.e., ungloved) hand. If an answer was given within 1.5 s, feedback (“correct” / “wrong”) appeared afterwards for 0.75 s; otherwise, a visual reminder to answer more quickly was shown.

Each participant’s recording session consisted of eight experimental blocks, with 32 trials each (balanced across task conditions within each block), resulting in 256 trials total. Depending on the duration of hand positioning, the delay period and occurrence of catch trials, each trial lasted between 8 and 15 s. On average, the main experimental session took about 60-65 min. To familiarize themselves with all conditions and cue meanings, participants performed a prior training session before the start of the recorded sessions, until they felt confident to complete the task. Between experimental blocks, participants could rest as much as they needed.

### Data preprocessing and analysis

Preprocessing was performed in MATLAB (MathWorks, version 2023b), with the *Edf2Mat* Matlab Tool (designed and developed by Adrian Etter and Marc Biedermann at the University of Zurich) used to for the conversion of EyeLink 1000 edf files; all statistical analyses steps were performed via MATLAB and JASP (Version 0.95.4). To derive hand movement data, finger flexion was recorded from seven sensors (two from each finger, one excluded due to technical difficulties) on a data glove (5DT Data Glove 14 Ultra MRI, 60 Hz sampling rate, full-speed USB 1.1 connection). Recorded data were averaged across all sensors and translated to a virtual hand position based on each participant’s individual glove calibration (see above). For analysis of gaze-hand tracking accuracy, hand movement data were interpolated to 1000 Hz to match the eye tracker’s recording frequency.

Analysis of the pupillometry data followed common guidelines (Steinhauer et al., 2022). The continuous pupil size measurements were segmented to select two relevant time windows per trial, the delay period (0 to 3000 ms post cue appearance) and the execution period 0 to 2000 ms post virtual hand appearance). As our analysis focused on pupil size changes, all values were then re-adjusted to a trial-specific baseline, defined as the mean pupil size during the 200 ms before the respective time window. Via a recorded reference of defined size, pupil size measures were converted from arbitrary units to square millimeters.

Gaze- and hand position data were similarly segmented (with the execution time window also including the 500 ms before the instructed movement onset, i.e. virtual hand appearance, to also include the ‘Ready’ signal), separated into x- (horizontal) and y- (vertical) direction, and re-adjusted with the exact midpoint of the screen as the zero baseline. Gaze- and virtual hand position values were then converted from position on the screen to degrees of visual angle. For each trial, the gaze-hand tracking accuracy time course was calculated as the offset between position of the visual target (i.e., the red dot on the tip of the virtual hand’s middle finger) and gaze position. Overall accuracy per trial was calculated by the root mean square error (RMSE) of this offset. Specific hand- and eye movements during the execution phase were detected, defined as any time window where hand- or eye movement velocity exceeded a threshold of 20% of total hand movement range per second (or 5 degrees visual angle per second, respectively). For each extracted movement, its onset, duration, and amplitude were calculated, as well as the number of saccades and mean saccade amplitude per trial.

Missing pupil size or gaze position data (e.g. due to blinks) was detected with the automatic algorithms of the EyeLink system. Data from trials with a continuous period of >500 ms or more than 20% (preregistered: 40%) of the entire time window missing were automatically excluded from our analysis. In minor cases of missing data, values were substituted via cubic interpolation from data 50 ms before and after the missing section. In addition to this automated algorithm, all raw data were visualized and inspected manually to identify possible technical issues or artifacts not recognized otherwise, with affected trials corrected or discarded. Across participants, 99.0% of trials were found valid for the delay period (SD: 1.4 %) and 98.6 % for execution period (SD: 2.4 %). Importantly, the valid trial rate of all participants was markedly above the threshold for exclusion defined in pre-registration (70% valid trials), with the lowest valid rates at 95% and 87 %, respectively. Therefore, no participant had to be excluded due to poor or missing data.

For pupil size data during the delay phase, the stimulus-locked mean time course and standard deviation were calculated on a single subject level for each condition, i.e., combination of factors *predicted visuomotor mapping* (congruent, incongruent), and *movement type cueing* (cued, uncued), averaging across up to 64 trials each. As an overall measure of pupil size change within the delay phase, we also calculated the condition-specific mean pupil size from 500 to 3000 ms post appearance of the visual cue for each participant, then used the results to perform a group-level repeated-measures analysis of variance (rmANOVA) with factors *visuomotor mapping* and *movement type cueing*. To determine the exact periods of significant differences between the (averaged) pupil size time series data of different conditions, we employed a nonparametric permutation cluster-based analysis (Maris & Oostenveld, 2007), implemented manually within our analysis. Specifically, for each comparison of two condition averages, we calculated a cluster-based t-test statistic between both (t-threshold / z-critical value for single time points at 1.96 for a two-sided test) to derive its corresponding test statistic. This observed test statistic was compared to the test statistics of 10,000 permutations, calculated from the time series signal of randomly switched conditions within participants. The p-value of the actually observed cluster was then determined as the proportion of random t-test statistics with an equal or larger test statistic than the one we recorded.

As described in the pre-registration, we wanted to determine the maximal pupil change during the delay phase. During analysis, we observed a period of initial pupil constriction. Therefore, we chose to calculate a ‘valley-to-peak’ amplitude between the time points with the lowest and highest pupil size within the delay phase for each condition and participant. We calculated two separate amplitudes: For the first, we subtracted minimum pupil size in the ‘valley’ time window (range: 500 to 1200 ms post cue appearance) from the maximum pupil size in the ‘early peak’ window (range: 800 to 2000 ms post cue appearance). In the second time window, minimum pupil size in the ‘valley’ time window was instead subtracted from the maximum pupil size in the ‘late peak’, window (range: 2500 to 3000 ms). For each amplitude, the associated duration, i.e. distance between time points was also calculated, while ensuring that the peak time point always occurred after the valley time point). The results were analyzed via group level rmANOVAs with factors *visuomotor mapping* and *movement type cueing*.

Analysis of the pupil size during the execution phase was performed concordantly, by first calculating the stimulus-locked mean time course and standard deviation for each condition. This included the nested factor *expectation violation* (matched, violated) besides factors *predicted visuomotor mapping* and *movement type cuein*g, averaging across 48 trials (expectations matched) or 16 trials (expectations violated) per condition. To determine overall differences between pupil dilation during movement execution, a three-way rmANOVA based on these factors was calculated for the average pupil size during movement execution (time window: 500 to 2000 ms post virtual hand appearance). To verifying that hand movement types (hand opening or closing) did not significantly affect pupil size, we also calculated means separated by hand movement for additional analyses.

To test whether expectation violation trials evoked a “surprise response” characterized by greater pupil dilation, a permutation cluster-based analysis (parameters see above) was calculated to determine time periods of significant differences between trials with matched vs violated expectations. To evaluate whether expectation violation-evoked pupil size changes differed depending on mapping and cueing conditions, we calculated the pupillary “surprise response” by subtracting, for each combination of factors (CC, CI, UC, UI), the average time course of trials with matched violations from ones with expectation violation. The resulting group means were used in a cluster-based permutation analysis (see above), contrasting cued and uncued movement type or congruent and incongruent mapping, respectively.

To evaluate how gaze-hand tracking performance during movement execution differed depending on trial conditions, the average time course of tracking accuracy was calculated on a single subject level for each condition. Additionally, condition means were calculated for tracking accuracy root-mean square-error (RMSE), hand movement onset, gaze movement onset (both relative to start of execution phase and hand movement onset), number of saccades per trial, mean saccade amplitude, and mean gaze acceleration. For all of these measures, three-way rmANOVAs with the factors described above were performed. A cluster-based permutation analysis with the same parameters as described above was performed on the mean tracking accuracy time courses of trials with matched and violated expectations. Following up on significant main or interaction effects, we calculated post-hoc t-tests; these contrasts were Holm-adjusted for multiple comparisons.

Finally, in a pre-registered exploratory analysis, we tested if pupil dilation during the delay period correlated with tracking performance during movement execution. For each participant, a single trial-based correlation analysis (Pearson correlation) was performed between pupil size change in the preparation phase (mean pupil size 500-3000 ms post cue appearance) and tracking performance, defined as either gaze-hand-tracking RMSE, onset of the first eye movement, or the number of saccades during movement execution. To test for significance on a group level, a one-sample t-test of the subjects’ correlation coefficients was performed, with the significance level Bonferroni-corrected for multiple comparisons.

## Results

### Task Compliance & Real Hand Movement

Participants showed excellent task compliance, with, on average, 99.1 % correctly executed hand movements (each participant > 94.1 %) and 98.3 % correct answers to the catch trials probing the cued visuomotor mapping (each participant > 89.8 %). On average, hand movements were initiated 306 ± 132 ms after the appearance of the virtual hand model and lasted for 445 ± 110 ms (Fig. 2). A three-way rmANOVA revealed a significant effect of movement type cueing on movement onset (*F*_(1,38)_ = 33.809, *p* = < .001, η²_p_ = .471), with pre-cued movements initiated on average 55 ms earlier than uncued ones. There was also a significant effect of predicted visuomotor mapping (*F*_(1,38)_ = 16.123, *p* = < .001, η²_p_ = .298), with participants, on average, initiating hand movements 25 ms later when expecting incongruent visual feedback. There was also a significant interaction effect of cueing and mapping (*F*_(1,38)_ = 23.499, *p* = < .001, η²_p_ = .382), with movement initiation especially delayed for uncued trials where participants expected incongruent feedback. We found no significant effect of expectation violation; but a significant interaction of expectation violation and visuomotor mapping (*F*_(1,38)_ = 9.339, *p* = .004, η²_p_ = .197). Hand movement durations were comparable across conditions and factors (no significant main effect or interaction).

**Figure 2:**
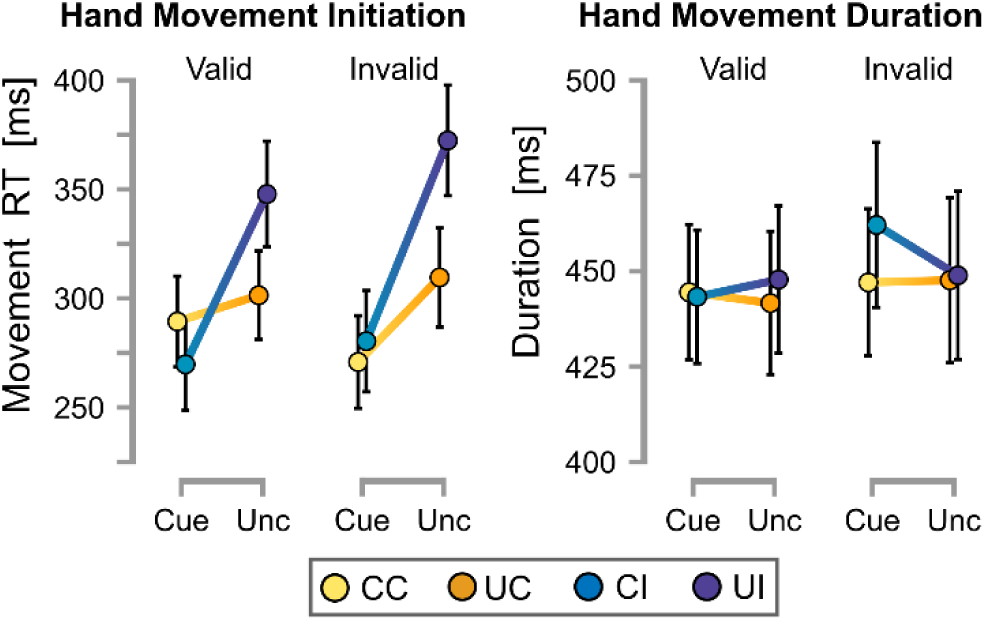
Hand movement execution. Mean hand movement initiation times after the appearance of the virtual hand (left) and mean duration of the hand movements (right). Error bars represents standard errors of the mean.

### Pupil Size Changes Related to Motor Planning and Visuomotor Prediction

We hypothesized that during movement preparation, the cue-induced pupil dilation should reflect the allocation of resources devoted to the generation of visuomotor predictions (incongruent>congruent, H1) and motor planning (cued>uncued, H2).

For the mean pupil size (500 to 3000 ms post cue appearance, see Table 2) a two-way rmANOVA revealed a significant effect of *movement type cueing* (*F*_(1,38)_ = 6.302, *p* = .016, η²_p_ = .142). Accordingly, the cluster-based permutation analysis of pupil size time courses (Fig. 3A) revealed a period of significant (*p* = .016) difference between cued and uncued trials from 718 to 2158 ms post cue appearance, with increased pupil dilation when movement type was cued in advance versus left ambiguous. The main effect of *visuomotor mapping* and the interaction effect were not significant.

**Table 2.** Pupil size change during the delay phase.

| Task conditions |  | Pupil size measures |  |  |
| --- | --- | --- | --- | --- |
| Mapping | Cueing | Mean size during delay phase | Valley-to-Early Peak amplitude | Valley-to-Late Peak amplitude |
| Congruent | Cued | $-0.772 \pm 0.625 \text{ mm}^2$ | $0.327 \pm 0.331 \text{ mm}^2$ | $0.237 \pm 0.581 \text{ mm}^2$ |
| | Uncued | $-0.882 \pm 0.630 \text{ mm}^2$ | $0.276 \pm 0.312 \text{ mm}^2$ | $0.217 \pm 0.547 \text{ mm}^2$ |
| Incongruent | Cued | $-0.686 \pm 0.584 \text{ mm}^2$ | $0.438 \pm 0.577 \text{ mm}^2$ | $0.362 \pm 0.751 \text{ mm}^2$ |
| | Uncued | $-0.815 \pm 0.594 \text{ mm}^2$ | $0.317 \pm 0.332 \text{ mm}^2$ | $0.378 \pm 0.513 \text{ mm}^2$ |

**Figure 3:**
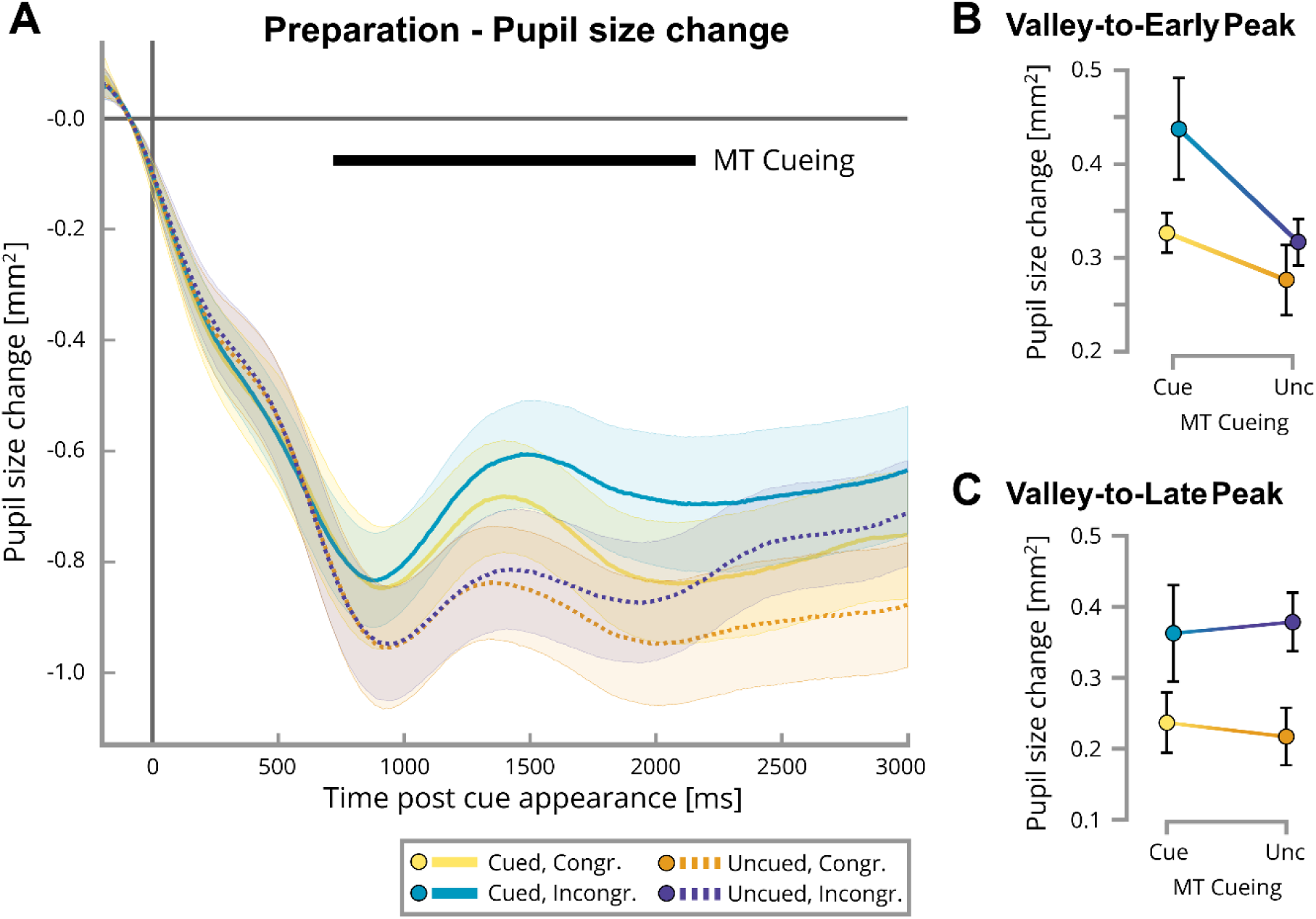
Preparation phase. **A)** Pupil size change in the delay period during the first three seconds after cue appearance, relative to baseline (200 ms before cue appearance). Each line displays one condition, i.e. combination of factors movement type (MT) cueing (cued, uncued) and predicted visuomotor mapping (congruent, incongruent), averaged across all subjects. Shaded areas indicate the standard error interval between subjects. The black bar indicates a cluster of time points showing a significant main effect of movement type cueing, with larger pupil dilation in cued > uncued trials (*p* < .05, determined via cluster-based permutation tests with n = 10,000). **B-C)** Group level means of the valley-to-peak pupil dilation analysis, i.e., the pupil size difference between the point of lowest pupil size (‘valley’, time window: 500-1200 ms) and the point of highest pupil size in either B) the ‘early peak’ time window (800–2000 ms) or C) the ‘late peak’ time window (2500–3000 ms). Error bars represent standard errors of the mean. For the early peak (B), the main effect of *movement type cueing* was significant (*p* = .030), with larger pupil dilation when the movement type was cued > uncued; the other main and interaction effect were not significant. For the late peak (C), the main effect of *visuomotor mapping* was significant (*p* = .024), with larger pupil dilation when incongruent > congruent mappings were cued; the other main and interaction effect were not significant.

Group level averages (Fig. 3A) showed both a common pupil dilation peak in the middle of the delay period, as well as an increase in pupil size towards the end of the phase. To test if task conditions affected pupil dilation differently during these two time periods, we calculated two separate valley-to-peak amplitudes, i.e. maximum pupil size changes (see Table 2; this separation was not pre-registered). For the valley-to-early-peak (Fig. 3B), a two-way rmANOVA confirmed a significant effect of movement type cueing (*F*_(1,38)_ = 5.109, *p* = .030, η²_p_ = .119), with a larger pupil dilation for cued>uncued trials, but no significant effect for predicted mapping or an interaction. For the valley-to-late-peak (Fig. 3C), in contrast, there was a significant effect of visuomotor mapping (*F*_(1,38)_ = 5.564, *p* = .024, η²_p_ = .128), with larger pupil dilation for trials where incongruent>congruent mappings were predicted, but no significant effect for movement type cueing or an interaction between both factors.

### Pupil Size Changes During Task Execution

We then tested for the hypothesized pupillary “surprise response”, reflected by increased pupil dilation during movement trials in which the visual movement feedback violated the cued visuomotor expectations (Hypothesis H4). As expected, a rmANOVA of the group mean pupil size 500 to 2000 ms post virtual hand appearance showed a clear, significant main effect of expectation violation (*F*_(1,38)_ = 65.774, *p* = < .001, η²_p_ = .634), with increased pupil dilation in response to violated>matched expectations (mean difference in pupil size = 0.47mm²). A cluster-based permutation analysis (Fig. 4A) revealed that the expectation violation effect was significant between 768 to 2002 ms after the appearance of the virtual hand (*p* < .001), i.e., largely after movement execution had finished.

**Figure 4:**
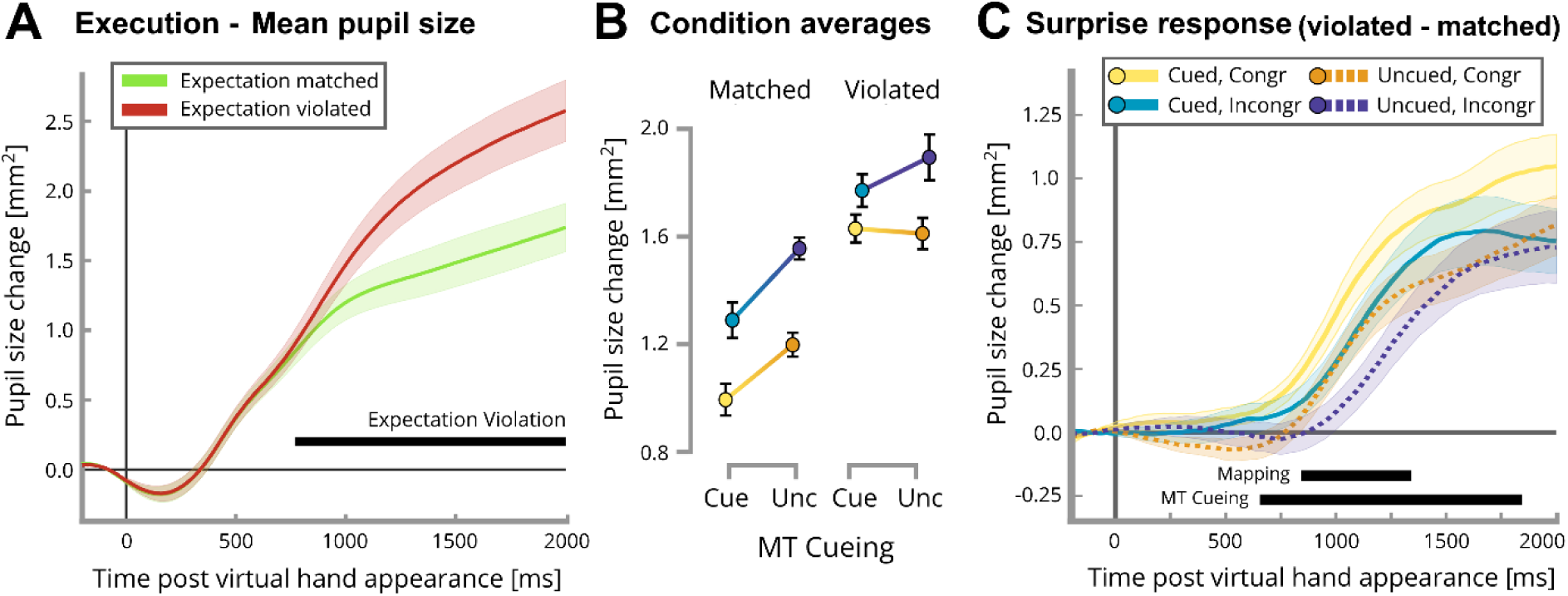
Pupil size during the execution phase. **A)** Pupil size changes during the execution of hand movements with visual hand movement feedback matching the cued expectations (green line) vs violating them (red line). Each line displays the group level average relative to baseline; 0 marks the onset of the execution period with appearance of the virtual hand. Shaded areas indicate the standard error interval between subjects. The black bar designates a significant cluster (*p* < .001, determined via cluster-based permutation tests with n = 10,000) of time points with a significant main effect of expectation violation. Pupil dilation was larger in trials where visuomotor mapping expectations were violated>matched. **B)** Group level mean pupil size during movement execution (500-2000 ms) relative to baseline; error bars indicate between-subject standard errors of the mean. In addition to the significant main effect of expectation violation, pupil dilation was significantly larger when incongruent>congruent mappings had been cued (*p* < .001), and when movement type had been left unspecified (uncued>cued, *p* = .002). Moreover, there was a significant interaction (*p* = .009) between movement type cueing and visuomotor mapping; the strongest expectation violation effect was observed in the cued, congruent condition. **C)** Average pupillary “surprise responses” per condition; i.e., the mean pupil size differences between trials were expectations were violated-matched. The black bars designate clusters of time points with a significant (*p* < .05) main effect of predicted visuomotor mapping (*p* = .036) and movement type cueing (*p* = .013), respectively. See Results for details.

We then calculated a three-way rmANOVA of pupil size during movement execution to test for possible conditional effects on pupil dilation (Fig. 4B). This revealed that cued visuomotor mapping had a significant effect on pupil diameter (*F*_(1,38)_ = 35.297, *p* = < .001, η²_p_ = .482), with relatively larger pupil dilation when participants expected incongruent mappings. The interaction of cued visuomotor mapping and expectation violation was not significant, but showed a statistical trend (*F*_(1,38)_ = 3.876, *p* = .056, η²_p_ = .093); i.e., the pupil dilation to expectation violation was somewhat, but not significantly stronger when congruent compared to incongruent visuomotor mappings had been cued. Movement type cueing also had a significant main effect (*F*_(1,38)_ = 11.525, *p* = .002, η²_p_ = .233), with relatively smaller pupil dilation when movement type was cued in advance. Here, the interaction between movement type cueing and expectation violation was significant (*F*_(1,38)_ = 7.521, *p* = .009, η²_p_ = .165), indicating a significantly different pupil dilation between invalid and valid trials (i.e., a stronger expectation violation effect or “surprise response”) when to-be-performed hand movements were pre-cued vs uncued. This interaction was driven by a particularly strong expectation violation effect in the CC condition (mean difference in pupil dilation between trials violating minus matching expectations = 0.64 mm²; all other conditions < 0.48 mm²), yet all individual expectation violation effects were significant (*p* < .003, corrected). All other interaction effects were non-significant.

For a time-resolved analysis of the above effects, we then calculated a cluster-based permutation analysis (Fig. 4C) on the condition time courses of the pupillary “surprise response”, i.e., grand averages of expectation violation minus expectation matched trials. This revealed a significant cluster (*p* = .013) for *movement type cueing* from 655 to 1842 ms after the virtual hand appeared, with a larger pupillary surprise response when movement type was cued>uncued. For *visuomotor mapping,* another cluster of significant difference (*p* = .036) was found from 814 to 1342 ms, with stronger pupillary surprise-related dilation when a congruent>incongruent visuomotor mapping had been cued.

### Gaze-hand Tracking

Next, we analyzed gaze-hand tracking performance to test for the hypothesized effects of expectation violation, visuomotor congruence, and movement cueing on execution (Hypotheses H4-H6).

First we tested for effects of expectation violation on hand-gaze tracking via a three-way rmANOVA. RSME (i.e., performance error) was significantly larger in trials where the visual movement feedback violated>matched the cued expectations (main effect, *F*_(1,38)_ = 29.522, *p* = < .001, η²_p_ = .437; Hypothesis 4). A cluster-based permutation analysis revealed that this performance difference was significant (*p* < .001) at 260 to 2000 ms after the appearance of the virtual hand (Fig. 5A,B). Furthermore, there was a significant interaction of expectation violation with movement type cueing on average gaze-hand-tracking RMSE (*F*_(1,38)_ = 8.716, *p* = .005, η²_p_ = .187, Fig. 5B); i.e., expectation violation more strongly impoverished gaze-hand tracking when movements were cued>uncued. There also was a significant interaction of expectation violation with cued visuomotor mapping (*F*_(1,38)_ = 7.768, *p* = .008, η²_p_ = .170); i.e., expectation violation more strongly impoverished gaze-hand tracking when congruent>incongruent mappings had been expected. Post-hoc contrasts revealed that the expectation violation effect (i.e., poorer gaze-hand tracking in trials violating than those matching cued visuomotor expectations) was significant whenever subjects expected congruent mappings, with the strongest difference in the cued, congruent condition (CC: *t*_(38)_ = 5.812, *p* < .001; UC: *t*_(38)_ = 5.250, *p* < .001). In contrast, there was no expectation violation effect under expected incongruent mappings (CI: *t*_(38)_ = 2.442, *p* = .232; UI: *t*_(38)_ = 0.457, *p* = 1). Furthermore, there was a statistical trend for a three-way interaction among the above factors (*F*_(1,38)_ = 3.615, *p* = .065, η²_p_ = .087).

**Figure 5:**
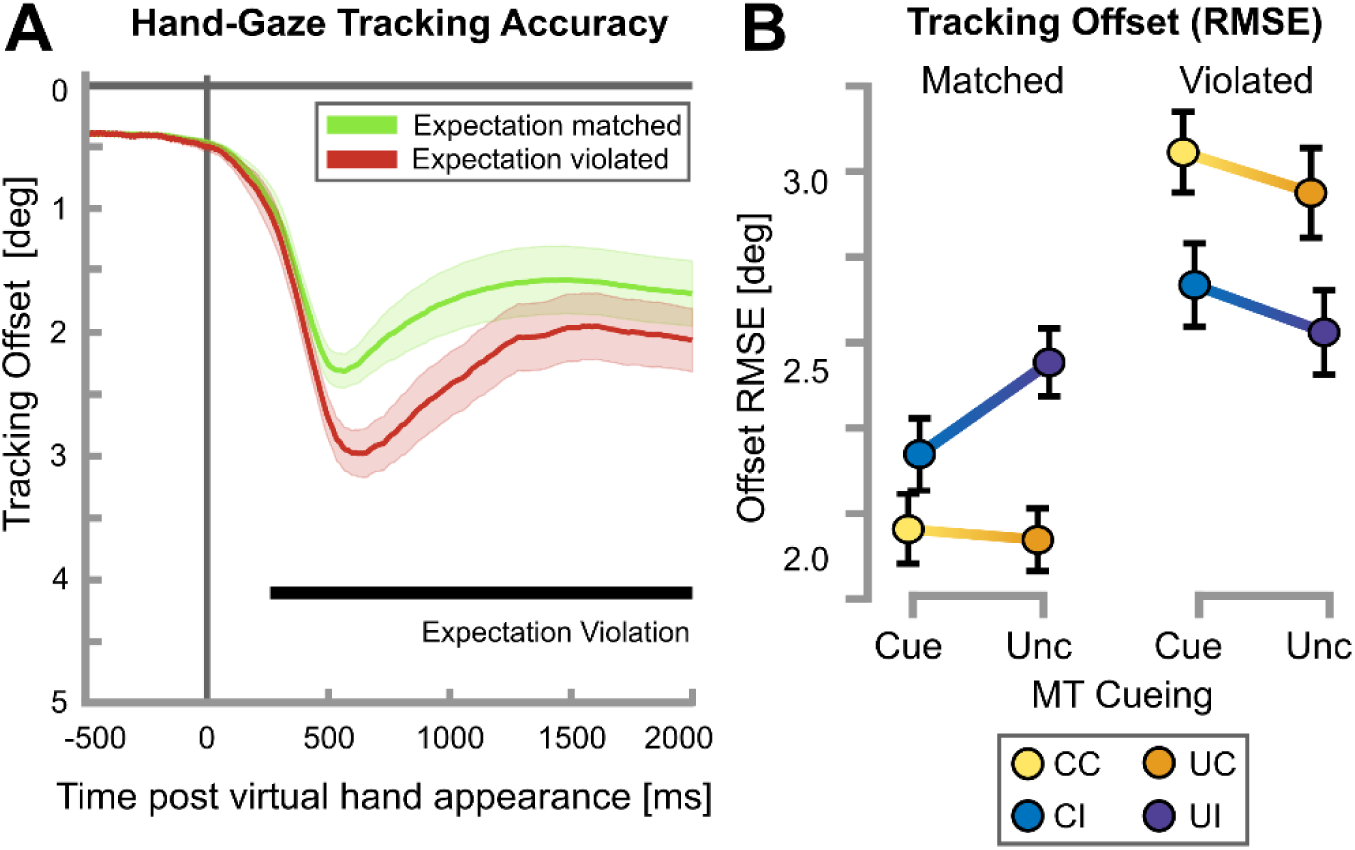
Gaze-hand Tracking Performance. **A)** Mean gaze-hand tracking offset during movement execution; i.e., the mismatch between the visual target’s position (red dot at the tip of the virtual hand’s middle finger) and the participants’ gaze position on the screen. Shown are the mean offsets of trials where the hand (and target) moved in accordance with cued visuomotor expectations (matched, green line) vs where it moved contrary to them (i.e., violated expectations, red line). The colored areas indicate between-subject standard errors of the mean. The black bar designates the cluster of time points with a significant main effect of expectation violation (*p* < .05, determined via cluster-based permutation tests with n = 10,000). I.e., gaze-hand tracking suffered from expectation violation. **B)** Group means of gaze-hand tracking offset (root mean square error within the time window 500 to 2000 ms post appearance of the virtual hand, see Methods); error bars indicate between-subject standard errors of the mean. Tracking offset was significantly larger when expectations were violated than when they were matched (*p* < .001). Moreover, this expectation violation effect interacted with predicted visuomotor mapping (*p* = .008) and movement type cueing (*p* = .005). See Results for details.

To understand this interaction and isolate the anticipated effects of movement type cueing and visuomotor mapping cueing on gaze-hand tracking performance, we analyzed trials with matched or violated expectations in two separate two-way ANOVAs. For trials with valid cueing, as expected, we found a significant effect of valid cueing (*F*_(1,38)_ = 8.071, *p* = .007, η²_p_ = .295), with better gaze-hand tracking performance under congruent compared to incongruent visuomotor mappings (*t*_(38)_ = 2.841, *p* = .022, Hypothesis 5). We also found an effect of movement type cueing (*F*_(1,38)_ = 15.915, *p* < .001, η²_p_ = .295), with better performance when the movement type was pre-cued (*t*_(38)_ = 3.989, *p* < .001, Hypothesis 6), and a significant interaction of cueing and mapping (*F*_(1,38)_ = 22.599, *p* < .001, η²_p_ = .373).

Post-hoc tests revealed that this effect was driven by a cueing advantage only for incongruent mappings: Here, when movement type was pre-cued, the subsequent gaze-hand tracking offset was significantly reduced compared to when it was left uncued (*t*_(38)_ = 5.418, *p* < .001). For congruent visuomotor mapping trials, in contrast, there was no such cueing advantage, with was observed for congruent visuomotor mapping trials, where tracking offset was similarly low both for cued and uncued trials (*t*_(38)_ = 0.865, *p* = 1). This difference in cueing advantage implied that, in cued trials, there was no performance difference between moving under congruent vs incongruent feedback (*t*_(38)_ = 1.608, *p* = .813).

The analysis of further eye movement kinematic parameters (saccade onset and amplitude, and tracking smoothness as indicates via gaze acceleration) overall supported these results; i.e., partly showing corresponding effects: Again, we found decreased tracking performance (later saccade initiation, larger and more abrupt saccades) for expectation violation, better performance under validly cued standard mappings, and earlier saccade initiation when movements were pre-cued (see Supplementary Fig. S1). Finally, pupil dilation during the delay phase did not significantly correlate with better tracking performance during movement execution.

## Discussion

Here, we used a VR-based, delayed-movement task with concurrent eye tracking to examine the oculomotor correlates associated with planning movements under standard vs nonstandard visual hand movement feedback (i.e., updating of visuomotor expectations); and violation of said cued expectations during execution. Our key findings were: During preparation, pupil diameter increased when the movement was pre-cued (vs left unspecified), and when nonstandard visual movement feedback was expected. Expectation violation impaired tracking performance and evoked a strong pupil dilation, which was strongest when participants had prepared specific (cued) movements and expected standard visuomotor mappings. Finally, gaze-hand tracking performance decreased under nonstandard mappings, but significantly less so when the to-be-executed movement was pre-cued. We now discuss these findings in more detail.

Firstly, as predicted (H1), we observed larger pupil dilation when participants expected incongruent compared to congruent visual movement feedback. This difference was observed during the later parts of the pupillary cue response (around 1 to 2 s post cue appearance) and became significant at the late peak. There are several possible interpretations of this pupillary effect. For one, cueing incongruent (nonstandard) visual hand position feedback may have led to the updating of internal visuomotor associations; i.e., the sensory predictions of the internal forward model. These updates would require increased computational resources and could consequently be associated with increased mental effort, which is known to evoke pupil dilation (see Introduction). This interpretation would also be supported by studies linking pupil dilation to error-based motor learning (e.g.,Yokoi & Weiler, 2022). However, pupil dilation may also have been evoked by the expectation of sensory conflict and a more challenging hand-eye tracking task. Higher recruitment of cognitive control reliably evokes increased pupil dilation Ariel & Castel, 2014; Rondeel et al., 2015; cf. van der Wel & van Steenbergen, 2018), and cue-evoked expectations of response conflict alone can evoke a larger preparatory pupil response (Unsworth et al., 2023). More generally, pupil dilation prior to task performance often reflects expectations about the upcoming task difficulty (Irons et al., 2017; O’Shea & Moran, 2016; White & French, 2016). While our current study does not allow to differentiate between these two interpretations, they need not be exclusive and may well be related. In sum, we propose the delay-period, visuomotor mapping-related pupillary responses likely reflected the mental effort associated with increased demand for cognitive resources when anticipating visual hand movement feedback conflicting with life-long visuomotor expectations; potentially, also related to updating visual forward predictions.

Secondly, also as predicted (H2), pupil dilation significantly increased after cues specifying the to-be-executed movements during preparation, compared to when the cues left the movement ambiguous until execution. We propose that this indicates the recruitment of cognitive resources by motor planning (Rosenbaum, 1980). Note that we observed significantly faster hand movement initiation in cued compared to uncued conditions; demonstrating a reaction time advantage of pre-cueing movements. This suggests that, in our task, participants prepared a specific movement if cued correspondingly, but abstained from movement planning in the uncued condition until the respective movement-specifying cue appeared in the execution period. It has been argued that the presentation of multiple response options may also, depending on context, lead to parallel motor planning (Cisek & Kalaska, 2002, 2005; Klaes et al., 2011; but see Dekleva et al., 2018). However, if participants would have planned both possible movements in parallel, one would have expected larger pupil dilation in the uncued condition, and less of a reaction time difference between cued and uncued conditions. Similarly, the pattern of our results suggests that participants did not perceive the uncued delay period as a state of increased behavioral (task-related) uncertainty, which one would expect to increase pupil dilation (Becker et al., 2024; Brunyé & Gardony, 2017; Fan et al., 2023; Preuschoff, 2011). In line with our interpretation, pupil dilation has previously been related to cues limiting the range of possible motor responses after instructed delays (Adam et al., 2014; Moresi et al., 2008). A similar effect was reported in two studies examining re-orientation of spatial attention, where directional cues predicting target position evoked larger pupil dilation compared to non-predictive cues (Dragone et al., 2018; Lasaponara et al., 2019). Not last, our findings match those obtained from our brain imaging study with a similar task design (Quirmbach & Limanowski, 2024), in which cortical motor system activity and reaction times suggested that if movements were left ambiguous, planning was reduced or delayed until execution. In sum, the most likely explanation of the cue-related pupil dilation is that it reflected the recruitment of cognitive resources by motor planning. While pupil dilation was most pronounced when specific movements were cued and visuomotor incongruence was expected, the anticipated interaction effect did not reach statistical significance (refuting H3). The timing difference between the significant main effects of movement type cueing and visuomotor mapping cueing on pupil dilation may indicate that motor planning and the preparation for novel visuomotor mappings constitute two sequential, potentially independent processes.

Thirdly, we observed significant “surprise responses” when, during execution, the visual movement feedback violated the cued expectations; i.e., when the virtual hand moved unlike predicted. These responses manifested themselves as increased pupil dilation, in line with our hypotheses (H4). Importantly, besides the significant main effect of expectation violation across conditions, there were conditional differences in the pupillary surprise response: Pupil dilation was significantly larger when the movement was pre-specified (cued>uncued), and when congruent compared to incongruent visuomotor mappings were expected. A significant interaction revealed that the pupillary surprise response was stronger after having planned a specific (i.e., cued>uncued) movement under expected standard (i.e., congruent>incongruent) visuomotor associations. The strength of expectations has been observed to modulate surprise-evoked pupil dilation (Kloosterman et al., 2015; Preuschoff, 2011; Reisenzein et al., 2019; Satterthwaite, 2007; Yokoi & Weiler, 2022a), and accordingly, a higher unexpectedness of stimuli produces a greater pupil dilation (e.g. Becker et al., 2024; Bianco, 2020; Preuschoff, 2011). In this light, our result suggests that when participants knew which movement they had to execute (cued>uncued), they could form stronger sensory expectations. Thereby, predictions of “standard” (congruent) visual movement feedback could have been stronger (more precise), because those associations (and forward models) had been learned and used life-long. Yet crucially, we observed a significant pupillary surprise response even when nonstandard (incongruent) visuomotor mappings had been cued (but movements were accompanied by standard visual feedback). Overall, this result shows that predictions of visual movement feedback can be adjusted prior to execution; i.e., in the absence of movement and related sensory inputs, demonstrating that forward models can be updated during motor planning, not only by sensory prediction errors (cf. Kuang et al., 2015).

Expectation violation also affected gaze-hand tracking; i.e., as predicted (Hypothesis H4), tracking was overall poorer when the visual movement feedback violated the cued expectations. Indeed, the effect of expectation violation was strong enough that when participants had to perform tracking under standard mappings, but nonstandard mappings were cued (and then violated), their performance decreased significantly in an otherwise simple tracking task. However, this effect did not reach significance when participants expected incongruent mappings. This could mean that any anticipatory processes during the preparation for movements under expected visuomotor incongruence (indicated by significantly stronger pupillary responses during the delay, see above) did not directly influence eye-hand coordination during subsequent movement execution. Alternatively, it could mean that even when unexpected, standard (congruent) visual hand movements were comparably easier to gaze-track, resulting in a relatively smaller error during those unpredicted (surprise) trials. This needs to be answered by future work. Nevertheless, surprise responses as quantified by pupil dilation and gaze-hand tracking showed, in principle, a consistent picture: In line with the surprise effect assessed via pupil dilation, the strongest detrimental expectation violation effect on tracking performance was observed in the cued, congruent condition. Furthermore, in the uncued, incongruent condition, there was virtually no expectation effect on tracking, and only a comparatively small expectation violation effect on pupil size.

Furthermore, gaze-hand tracking performance in valid trials was overall poorer under nonstandard mappings; which confirmed our Hypothesis H5 and mirrored previous hand-eye coordination findings (Chen et al., 2016a; Dalecki et al., 2019; Hosseini, 2017; Landelle et al., 2016; Yeomans et al., 2021). Crucially, when the movement type was pre-cued, this difference was significantly reduced, and not significantly different from that under standard visuomotor mappings. This suggests that, during motor planning, forward predictions of visual movement feedback could be successfully updated—benefiting subsequent eye-hand coordination under nonstandard visuomotor mappings (cf. Chen et al., 2016b; Danion & Flanagan, 2018; Landelle et al., 2016; Matsumiya, 2021; Vercher et al., 1995; M. A. Yeomans et al., 2021). No significant cueing-evoked difference was found for tracking under congruent feedback, which could suggest that for a standard mapping (i.e., following life-long learned associations), motor planning may have been easier or occurred faster, so that pre-cueing provided no behavioral benefit. Thus, while a cueing benefit on eye-hand coordination was not observed under congruent (standard) visuomotor mappings, the significant cueing benefit under nonstandard mappings partly supports our Hypothesis H6.

Our results should be interpreted in light of the following limitations. Unexpectedly, participants exhibited pupil constriction during the first 900 ms of the delay period; which could have been caused by changes in luminance and/or residual cognitive processing during pre-positioning of the hand based on visual feedback before each trial. Furthermore, we cannot conclusively determine to what extent eye movements in our task actually represented intended hand tracking or simply the parallel planning and execution of hand and (anti-)saccade movements. Further research using prolonged and more complex hand movements could clarify this distinction.

In conclusion, our results demonstrate that oculomotor responses encode processes related to motor planning and flexible forward prediction of sensory action consequences ahead of execution; and that these predictive processes affect eye-hand coordination and pupillary responses during subsequent execution.

## Acknowledgments

This work was funded by the German Research Foundation (DFG, Deutsche Forschungsgemeinschaft) as part of Germany’s Excellence Strategy – EXC 2050/2 – Project ID 390696704 – Cluster of Excellence “Centre for Tactile Internet with Human-in-the-Loop” (CeTI) of TUD Dresden University of Technology. JL was supported by a Freigeist Fellowship of the VolkswagenStiftung (AZ 97-932). We thank Gesche Vigh for help with implementing the virtual reality.

## Disclosure statement

The authors declare no conflict of interest.

## Data availability statement

The behavioral and eye tracking data will be made available upon request.

## Notes

### Competing Interest Statement

The authors have declared no competing interest.

