## Supplementary Material for "Eye tracking insights into movement preparation and execution under nonstandard visual movement feedback"

### Fig. S1: Secondary hand-gaze tracking performance indicators

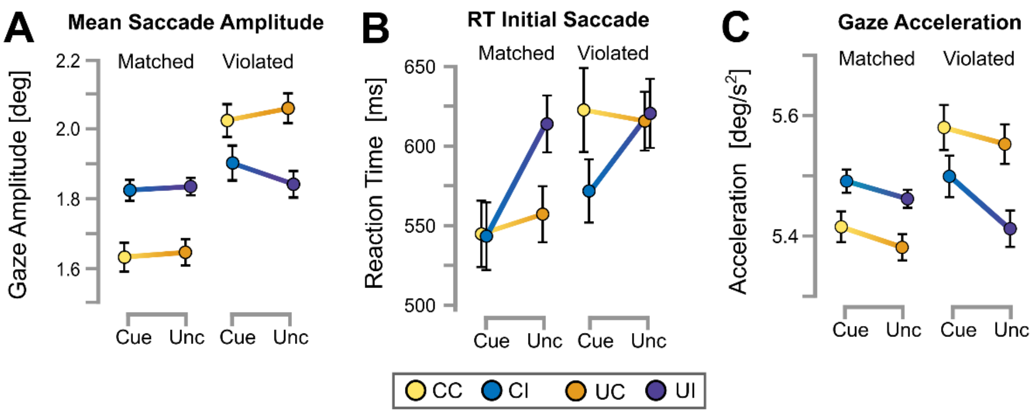


Group means of secondary indicators of hand-gaze tracking performance, i.e., (A) saccade amplitude during movement execution, (B) reaction time until initiation of the first tracking saccade, and (C) gaze acceleration during the execution phase; error bars indicate between-subject standard error of the mean. Analysis via three-way rmANOVAs revealed a significant effect of expectation violation for all measures, with violated expectations associated with increased saccade amplitude ($F_{(1,38)}$ = 24.964, $p$ = < .001, $\eta^2_p$ = .396), later saccade initiation ($F_{(1,38)}$ = 20.666, $p$ = < .001, $\eta^2_p$ = .352), and higher gaze acceleration, i.e. more abrupt eye movements ($F_{(1,38)}$ = 7.918, $p$ = .008, $\eta^2_p$ = .172). Expectation violation furthermore interacted with predicted visuomotor mapping ($p$ = < .001, for all measures), as the performance decrease following violated expectations was especially pronounced when participants expected standard visuomotor mappings. For saccade initiation (B), there was also a significant main effect of movement type cueing ($F_{(1,38)}$ = 12.222, $p$ = .001, $\eta^2_p$ = .243), with earlier initiation when movements were pre-cued, and an interaction of cueing with predicted visuomotor mapping ($F_{(1,38)}$ = 18.761, $p$ = < .001, $\eta^2_p$ = .331), as this cueing advantage was strong in trials under incongruent mapping, but not significant for congruent mappings. We also found a (non-significant) trend for an interaction of cueing and expectation violation ($F$ = 3.347, $p$ = .075, $\eta^2_p$ = .081). For gaze acceleration (C), there was also a significant effect for cueing ($F$ = 5.002, $p$ = .031, $\eta^2_p$ = .116), with higher acceleration (i.e., more abrupt saccades) when hand movements were pre-cued.
